# Effects of Alzheimer’s disease plasma marker levels on multilayer centrality in healthy individuals

**DOI:** 10.1101/2024.01.08.573504

**Authors:** Alejandra García-Colomo, David López-Sanz, Ignacio Taguas, Martín Carrasco-Gómez, Carlos Spuch, María Comis-Tuche, Fernando Maestú

## Abstract

Finding early and non-invasive biomarkers that help identify individuals in the earliest stages of the Alzheimer’s disease continuum is paramount. Electrophysiology and plasma biomarkers are great candidates in this pursuit. Furthermore, the combination of functional connectivity metrics with graph-theory analyses allows for a deeper understanding of network alterations. Despite this, this is the first MEG study to assess multilayer centrality considering inter-band connectivity in an unimpaired population at high risk of Alzheimer’s disease. Our objective is twofold. First, to address the relationship between a compound centrality score designed to overcome previous inconsistencies stemming from the use of various individual metrics, and plasma pathology markers of Alzheimer’s disease in unimpaired individuals with elevated levels of the latter. Lastly, to evaluate whether hubs’ centrality is more affected by the pathology.

33 individuals with available MEG recordings and elevated plasma pathology markers were included. A compound centrality score for each brain source of every subject was calculated combining widely used centrality metrics, considering intra- and inter-band connections.

Spearman correlations were carried out to address the association between each node’s centrality score and biomarkers levels. Next, to test whether greater associations were found in hubs, a correlation between the obtained rho and the grand-average of the centrality score was carried out.

Increasing concentrations of p-tau231 were associated with greater centrality within the network of posterior areas, which increased their connectedness in the theta range with the remaining areas, regardless of the latter’s frequency range. The opposite relationship was found for left areas, that decreased their connectedness in the gamma frequency range. Hubs’ centrality was significantly more affected by p-tau231 levels.

Our results expand previous literature demonstrating early network reorganizations associated with elevated plasma p-tau231 in cognitively unimpaired individuals. Multilayer centrality increases in the theta band in posterior regions are congruent with previous results and theoretical models, that predict a longitudinal evolution towards a loss of centrality. On the other hand, the changes in multilayer centrality found in the gamma band could be associated with inhibitory neuron dysfunction, classical in AD pathology. Lastly, hubs were more likely to increase their centrality in association to p-tau231, thus corroborating hubs’ vulnerability.

## Introduction

Alzheimer’s disease preclinical phase is characterised by the appearance of neuropathological changes with no associated clinical symptom^1^ and begins up to 20 years before dementia onset. This highlights the importance of finding early and non-invasive biomarkers.

With the emergence of the first pathological hallmarks, neuronal dysfunction becomes manifest in the form of functional changes.^2,3^ Electrophysiological recordings of brain activity of individuals at the different stages of the continuum reveal a progressive disruption of functional connectivity (FC), marked by an initial increase in connectivity followed by a state of hypoconnectivity, first observed in posterior regions.^4–7^ This inverted u-shaped pattern is thought to be, at least partly, related to amyloid beta (Aβ) accumulation^8–10^ due to the positive feedback loop between Aβ and hyperexcitation and hyperconnectivity.^10–12^ Given this early relationship between neuronal activity and Aβ, areas with higher neuronal activity might be at a greater risk of dysfunction as reflected by their early affectedness in the continuum.^8^

In the context of network neuroscience, that defines the brain as a graph, centrality measures represent the nodes’ importance for the functioning of the network on local, global, and intermediate scales.^13,14^ Regarding the prodromal and dementia phases, a progressive loss of centrality in posterior areas and an increase in anterior ones, affecting predominantly association areas, in a disease severity dependent fashion has been described.^15–19^ In the preclinical phase, however, divergent results emerge, potentially due to the use of different scale centrality metrics.^20,21^ Studies performed using degree centrality report a positive association with Aβ-PET levels along with an increase in hubness in temporo-occipital,^10,22^ whereas eigenvector centrality studies report reductions associated with Aβ pathology, affecting mostly posterior regions.^14,23^ Moreover, given the intrinsic complexity of electrophysiological signals, most studies tend to adopt a single-layer subnetwork approach that considers each frequency band independently, which in combination with the different centrality measures, sometimes yield inconsistent results. Duan et al.^24^ report centrality alterations, of different signs depending on the measure, in different frequency bands for mild cognitive impairment and Alzheimer’s disease patients. In an attempt to use a more integrative approach, Yu et al.^19^ show that, when combining centrality information from different bands, changes in hubs emerge that don’t appear when addressing single frequency band networks. This study, however, doesn’t consider interlayer links.

Given the limitations of PET and CSF markers that make them poor candidates for early detection and to track disease progression (i.e. high costs, invasiveness and low accessibility) emphasis has been placed on plasma biomarkers that have proven reliable as indicators of Alzheimer’s disease pathology.^25^ High levels of neurofilament light (NfL) chains are indicative of axonal injury and neurodegeneration. Although this is not a specific biomarker of Alzheimer’s disease some authors report significant increases early on in the continuum.^25^ The recently discovered species of phosphorylated tau at threonine 231 (p-tau231) presents abnormal levels, both in plasma and CSF, that can be detected as early on in the continuum as the preclinical phase. Its levels increase in parallel to and tracking Aβ increases even before PET positivity for this marker has been achieved.^26–30^

The present study is, to our knowledge, the first to address the relationship between NfL and p-tau231 plasma levels, with a multilayer compound centrality metric in cognitively unimpaired individuals who present elevated plasma pathology markers. The multilayer compound centrality score is based on functional connectivity measures from magnetoencephalography (MEG) recordings and the rationale behind it is twofold. Firstly, a multilayer approach will provide an integrative perspective of the network properties, while the development of a compound score will contribute to bridge the discrepancies found in the literature stemming from the use of different centrality measures. We expect to find an association between the plasma biomarkers’ levels and the centrality score of posterior regions. Additionally, we aimed to study how this relationship was affected by the relevance of a given region in the network. Given the association between neuronal activity and Aβ deposition, we expect to find that the centrality of regions with a hub-like behaviour in the network (i.e. areas of great relevance and high neuronal activity) will be particularly affected by Aβ as reflected by p-tau231 levels.

## Results

### Association between plasma markers and centrality score

Firstly, NfL levels’ association within each individual source’s centrality score were studied by means of correlation analyses. As a result of said analyses no significant clusters emerged in any of the bands tested. Henceforth, we did not find any significant association between centrality score and NfL levels.

Regarding p-tau231 levels and its association with centrality score, two clusters were found to be significant after multiple comparisons correction. In the theta band, we observed a significant cluster (rho_sum_ = 60.9; p_cluster_ = 0.039) in which p-tau231 plasma levels and the centrality score were positively correlated (Figure 1 A). This cluster included mostly bilateral regions covering posterior aspects of the brain. These included mainly areas over parietal, occipital and posterior temporal regions such as the angular gyrus, inferior parietal lobe, inferior and middle occipital cortices, fusiform gyri and left precuneus among others. Centrality score in the gamma band was also significantly associated with p-tau231 levels (rho_sum_ = -108.45; p_cluster_ = 0.016) although in a negative direction (Figure 1 B). Negative correlations between these two metrics were mainly observed over left hemispheric regions including a great proportion of the left temporal lobe (both medial and lateral structures) and extending into ventral and posterior aspects of the frontal and parietal lobe (such as the inferior frontal and posterior frontal gyri).

**Figure 1.**
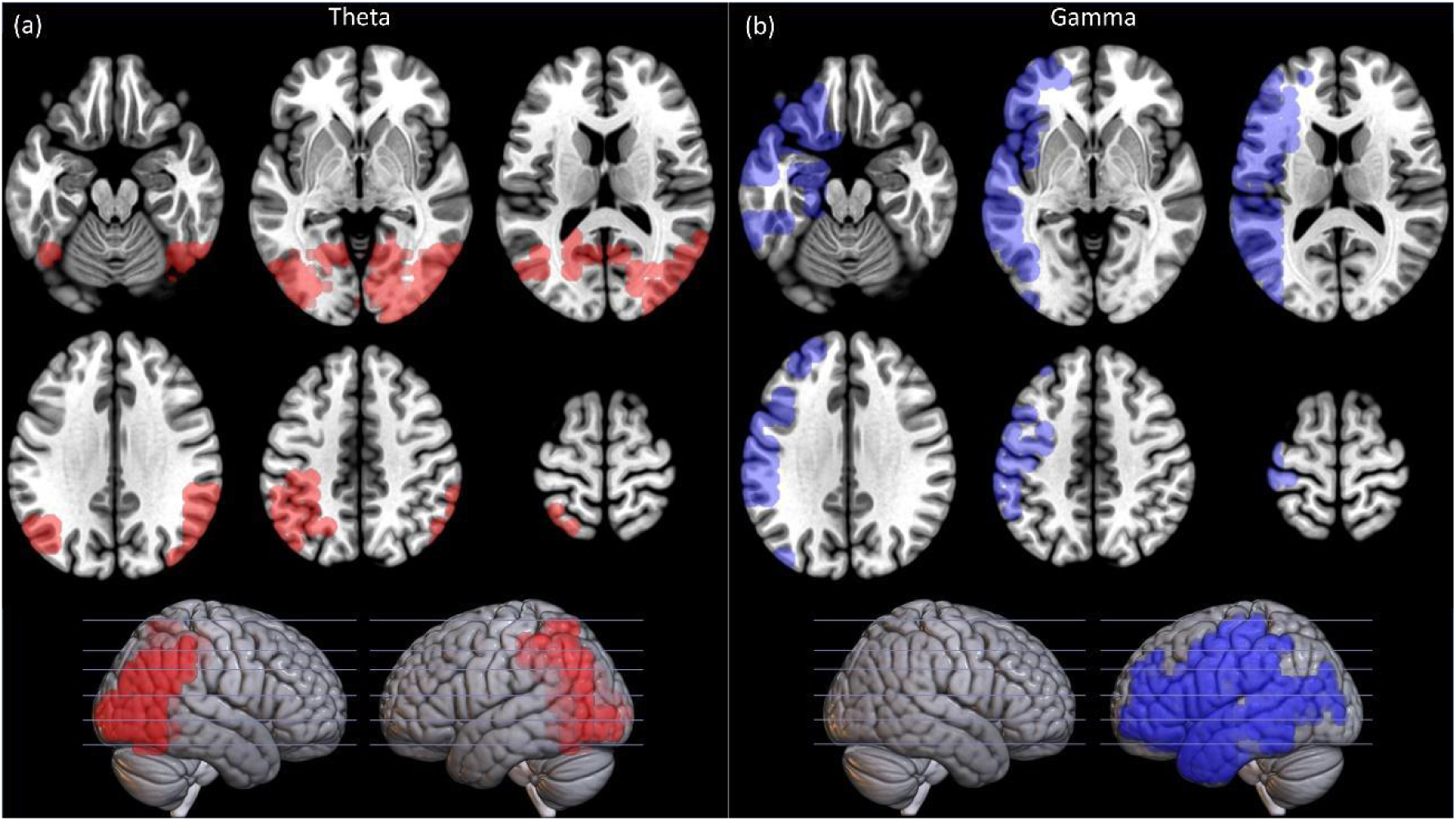
Regions showing significant correlation between the centrality score and p-tau231. **(a)** Regions that show a significant correlation with the theta band centrality and **(b)** regions that show a significant correlation with the gamma band centrality. Red colors denote positive correlation coefficients while blue colors indicate negative correlation coefficients.

### Grand-average centrality affects the association between plasma markers and centrality score

To this aim we conducted two correlations as previously explained, one for each band showing a significant association between the centrality score and the plasma marker. Consequently, we conducted the analysis to study how the mean relevance of each brain region in the network (i.e. its grand-average centrality score) influenced the relationship between p-tau231 levels and theta/gamma centrality score for each source.

Results for the theta band showed a highly significant association between the mean relevance of each brain region, and how their centrality changes in association with p-tau231 levels (rho = 0.499; p = 9.2 x 10^-77^). This positive association indicates that the more relevant a node is in the network, the larger is the association between its centrality in the theta band and p-tau231 level. This relationship can be further observed in Figure 2, where it can be appreciated how the association (rho values) between p-tau231 levels and theta band centrality is larger in those sources with a higher grand average centrality in the sample. Thus, increasing linearly with the general relevance of the node in the network.

**Figure 2.**
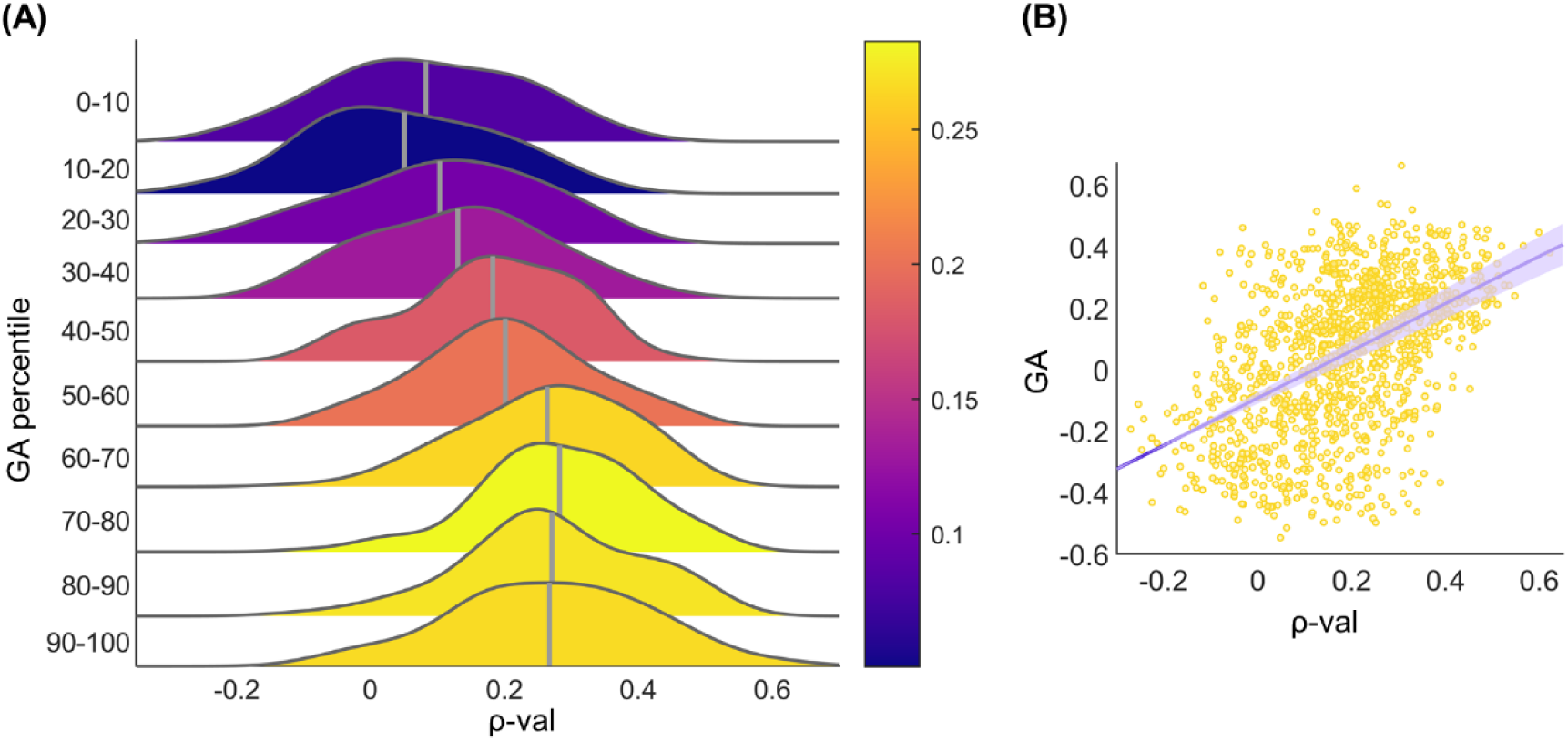
Influence of the total relevance of each node within the network (i.e. grand-average centrality) on the correlation between *centrality score* and *p-tau231 levels*. **(a)** Probability distribution for the correlation coefficients between *centrality* score and *p-tau231 levels* belonging to nodes of increasing relevance in the network, divided in increasing 10 percentile steps. Gray vertical lines indicate mean correlation coefficient (ρ-val) within that GA percentile step. **(b)** Scatter plot of the correlation coefficient (ρ-val) of all nodes as a function of their grand-average centrality.

In the gamma band, we did not observe a significant association between these two variables (rho = 0.006; p = 0.81).

## Discussion

This pioneer study is, to the best of our knowledge, the first effort to assess multilayer centrality considering interlayer links in an unimpaired population at high risk of developing dementia using MEG. Moreover, it is also the first one to address the relationship between network centrality alterations and the Alzheimer’s disease pathology plasma markers p-tau231 and NfL. Our results reveal a twofold relationship between p-tau231 concentrations and the functional network properties among individuals with higher levels of plasma Alzheimer’s disease biomarkers. With increasing concentrations of p-tau231, posterior areas hold greater relevance within the network through their theta oscillations in coordination with the activity of all other regions in all other frequency bands, given the multilayer nature of the networks studied. In contrast, left hemispheric regions show the opposite pattern, exhibiting a lower level of importance within the network through their activity in the gamma frequency band. Lastly, a subsequent analysis revealed that the most relevant areas for the network are also the ones that present the greatest association between their integrative centrality score and p-tau231.

In recent years, plasma p-tau231 has proven highly relevant for the study of the preclinical stage of Alzheimer’s disease given its early increases that parallel those of Aβ species. Indeed, it shows elevated levels even before Aβ-PET positivity is achieved, is a good predictor of elevated Aβand is discriminant of the different stages of the continuum.^26–30,40^ Consequently, in this study, p-tau231 has been considered a proxy for Aβ and Alzheimer’s disease pathology. Our results show heightened and reduced centrality in the theta and gamma frequency bands, respectively, in relation to plasma p-tau231 levels. Regarding the theta band, posterior areas that include the bilateral inferior parietal lobe, angular gyri, middle temporal gyri and inferior and middle occipital regions, become more relevant within the network by increasing their connectedness within the 4-8Hz range with the remaining areas, regardless of the latter’s frequency range, as p-tau231 levels increase. This is congruent with previous literature, where alterations in the theta frequency band have been extensively described. Across the continuum, progressive increases in relative theta power,^41,42^ along with enhanced synchrony in this frequency band have been consistently observed.^5,43–45^ Furthermore, Hatz et al^44^ found the theta connectivity between central and parieto-occipital regions to be the most discriminant characteristic among healthy controls, mild cognitive impairment, and Alzheimer’s disease patients. Finally, these posterior regions have been extensively studied in the context of Alzheimer’s disease, Aβ pathology, and display centrality alterations across the continuum.^14,15,19,23,46^ Interestingly, our results expand previous findings by demonstrating that not only local and long-range synchronization are affected, but also network structure is reconfigured this early in the Alzheimer’s disease continuum in association with novel Alzheimer’s disease plasma markers. This network reconfiguration is highly relevant and should be further studied as it could underlie the functional and cognitive changes observed in the Alzheimer’s disease continuum, thus providing a deeper understanding of the pathology.

On the other hand, network centrality from regions in the left hemisphere, with a predominance for the frontal and temporal cortex, exhibit a negative correlation with p-tau231 levels; the relevance of these areas within the network diminishes as gamma activity participation decreases in the overall network communication structure in relation to p-tau231 concentrations. Gamma activity has been extensively linked to parvalbumin inhibitory neurons; specifically, alterations in these neurons suppress gamma activity.^47–49^ On a related note, Verret et al^47^ demonstrated increases in hypersynchrony associated with gamma activity reductions in an experimental model of AD, Cuesta et al^50^ reported decreased gamma connectivity in association to epileptogenic activity, and various works yield promising findings regarding the use of invasive and non-invasive techniques to ameliorate of Alzheimer’s disease pathogenesis and symptomatology by promoting neural activity in this frequency band.^51,52^ Consequently, the diminished relevance of these areas within the network in the gamma frequency band could be a reflection of ongoing Aβ pathology and neuronal hyperactivity, commonly observed at the beginning of the disease.^9,10^

The lack of association between NfL, as opposed to p-tau231, with the centrality score might be due to different factors. On the one hand, it might be that these individuals are still in a very early stage of the Alzheimer’s disease continuum for NfL levels to be discriminant. In their respective studies, Mattson et al^53^ and Nyberg et al^54^ found significant differences in NfL levels in individuals further along the continuum, but failed to find them in preclinical individuals, and argue that this could be due to the minimal axonal injury found during this stage, the disease-inespecificity of this axonal-damage marker and/or the increases in this marker associated with age. Although some studies do report significant differences in NfL levels in the preclinical stage, their sample size is very extensive.^25^ Hence, the reduced number of subjects in the present study might represent a limiting factor.

Subsequently, to study whether hubs are more vulnerable, we calculated a grand-average measure that considers the relevance of each node in all frequency bands as the average across subjects to determine the importance of every node in the functional network. Our results are in agreement with previous studies, suggesting that those areas showing a more hub-like behavior are the ones more intensely affected by p-tau231 early on in the disease continuum. These regions showed the highest association between the increased centrality score and the p-tau231 plasma levels and include areas such as the bilateral inferior parietal gyrus, temporal regions, and occipital areas, which have been defined as hubs, as well as vulnerable to Alzheimer’s disease pathology, both in network studies as well as in classic FC studies.^4,10,19,55^

On the other hand, while we describe that the most relevant areas are the ones that exhibit a larger increase in centrality associated to the p-tau231 levels, previous literature in later stages of the continuum have described decreases in centrality in these same regions.^4,16,19^ This pattern of an initial increase that gives way to a decrease in centrality can be understood through the theoretical model proposed by Stam^17^, which explains why hubs exhibit an increased vulnerability in diseases like Alzheimer’s disease following this inverted U-shape centrality pattern. Accordingly, in the early stages, subordinate nodes in the hierarchy start sending information directly to hubs. This reorganization increases their centrality but, if maintained, produces hub overload, which makes them lose centrality in favor of other areas that increase their own. This pattern of posterior hub disruption accompanied by an anteriorization of the most central areas has been extensively described in later stages of the continuum.^4,16,19^ Moreover, it is congruent with computational models that demonstrate that hub vulnerability can be activity-dependent; excessive neural firing, which has been demonstrated in the early stages of AD, is associated with the increase in Aβ species and begins in some of these regions.^10,18,56^ This neural hyperexcitability induces synaptic dysfunction that would potentially translate into increased connectivity and centrality, that eventually give way to a reduction in connectivity, more random networks, and selective damage to hubs.^17,56^

Given the aforementioned treatment of p-tau231 as an Aβ proxy, the relationship between centrality and p-tau231, that is maximum in hubs in the theta frequency band, might indicate that these regions are starting to exhibit pathological changes on a molecular level that underlie the ones observed on a functional level, thus suggesting that these individuals are entering the Alzheimer’s disease continuum.

The importance of performing multilayer analyses to yield a more comprehensive understanding of brain function has already been discussed in the literature.^19^ Likewise, the use of an integrative centrality score allows the study of complementary aspects of the individual and classically studied measures, which, on their own, have been known to yield conflicting results.^20,21^

This is an unprecedented study aimed at providing an integrative understanding of brain network patterns in cognitively unimpaired individuals at risk of Alzheimer’s disease due to their concentrations of emerging plasma biomarkers. However, this study is not without shortcomings. As previously mentioned, the sample size could represent a limiting factor, preventing the finding of a relevant relationship between NfL levels and brain dynamics. Remarkably, this study is part of an initiative aimed at longitudinally following these individuals for decades, with the next time-point assessment already approved and plans for cohort enrichment. Additionally, future assessment rounds should include, in addition to plasma biomarkers, other well-established techniques to quantify Alzheimer’s disease pathology to allow for a better classification of individuals and reinforce the validity of these emerging ones. Nevertheless, plasma biomarkers might represent a more ecological, less invasive, more cost-efficient, and widely accessible alternative.

Overall, our study takes a pioneering approach to study functional networks in individuals with elevated levels of Alzheimer’s disease pathology markers, being the first to combine the use of MEG data, plasma biomarkers and cognitively unimpaired individuals, with novel and integrative multilayer network analyses. This allowed us to overcome the oversimplification of single-frequency band studies and combine the properties of various centrality measures to obtain a more comprehensive understanding of brain connectivity patterns. Our results are consistent and expand previous literature, showing early alterations in network communication associated with elevated plasma biomarkers in cognitively unimpaired individuals, particularly in hubs, which are vulnerable areas to Alzheimer’s disease pathology.

## Methods

### Participants

This study was carried out in a sample of 33 participants extracted from a group of 139 participants, all of them with available plasma markers and a valid MEG recording. Volunteers for this study are part of a larger project longitudinally following participants’ evolution over time in subsequent waves. This project aims to study the earliest electrophysiological signs of probable Alzheimer’s disease pathology by combining different neuroimaging, neuropsychological and plasma measurements for each participant at different stages. The data included in this work belongs to the second wave for which plasma markers (NfL and p-tau231) were acquired for the first time in their follow-up. Given our interest in studying the earliest signs of Alzheimer’s disease pathology in a preclinical sample, we selected only those participants exhibiting higher values of the two plasma markers available. To maximize the likelihood of detecting early Alzheimer’s disease signs, only participants with an above-median value for both markers were included. This means that participants included were those who were above the 50^th^ percentile in both NfL and p-tau231 values distributions.

All participants underwent a thorough neuropsychological assessment to ensure a healthy cognitive status in the different domains. Table 1 summarizes relevant demographic and clinical information of the sample.

**Table 1.** Sample Demographic information. Values are presented as Mean ± Standard Deviation.

|  | Mean ± std<br>(n=33) |
| --- | --- |
| Age | 61.8 ± 7.1 |
| Sex | 11/22 |
| MOCA | 27.3 ± 2.2 |
| NfL | 2.1 ± 1.9 |
| p-tau231 | 482.6 ± 138.3 |

Each participant had an available MRI scan acquired during the first wave (2 to 3 years prior to the second wave). During the second wave, all participants underwent an MEG scan and a blood sample extraction for plasma markers determination.

Exclusion criteria for the current study included: (1) history of psychiatric or neurological disorders or drug consumption in the last week that could affect MEG activity; (2) family history of dementia other than Alzheimer’s disease; (3) evidence of infection, infarction, or focal lesions in a T2-weighted MRI scan; (4) alcoholism or chronic use of anxiolytics, neuroleptics, narcotics, anticonvulsants, or sedative-hypnotics; (5) a score in the Montreal cognitive assessment below 26 and (6) unusable MEG recording or T1-weighted image.

### Plasma sample determinations

Plasma p-tau231 and NfL concentrations were quantified by competitive enzyme-linked immunosorbent assay, using commercial kits (MyBiosource, Inc., USA, MBS724296, and human NF-L Elabscience, respectively), according to the manufacturers’ instructions. Tests were performed in duplicate, and an automated microplate reader (Biochrom ASYS UVM 340, Cambridge, UK) measured the optical density at 450 nm with Mikrowin 2000 software (Berthold Technologies, Germany).

### MEG recordings and preprocessing

Four minutes of ongoing brain activity during resting-state under eyes closed condition were acquired from each participant at the Center for Biomedical Technology (Madrid, Spain) using a 306-sensor Vectorview MEG system (Elekta AB, Stockholm, Sweden). Continuous head position information during the recording was registered through four coils placed over the forehead bilaterally and both mastoids of each participant. To this aim, coil position and head shape information were digitized using a Fastrack Polhemus system (Polhemus, Colchester, VT, USA). Electrooculographic and electrocardiographic activity were registered by placing two sets of bipolar electrodes. The recordings were performed inside a magnetically shielded room, and participants were instructed to stay as still as possible and relax. Data was acquired at a sampling rate of 1000 Hz and filtered online between 0.1 and 330 Hz. Before preprocessing of the signal, the contribution of head movements and distant sources outside the brain were removed from the signal by using the spatiotemporal expansion of the signal space separation method,^31^ implemented by Neuromag Software (MaxFilter version 2.2, correlation 0.90, time window 10s).

After the spatiotemporal expansion of the signal space separation, MEG signal was preprocessed using Fieldtrip to detect muscular, ocular and jump artifacts.^32^ Ocular and cardiac components were removed from the signal by using an independent component analysis based algorithm. After data segmentation, only subjects with at least 20 valid segments of 4 consecutive seconds of artifact-free activity were kept for further analyses. Given the elevated redundancy of the data after applying the signal space separation method,^33^ only data from magnetometers were considered for subsequent analyses.

### Source reconstruction (individual MRI)

Individual T1 structural brain images were used to reconstruct source space MEG signals. To do so, MRI was acquired in a General Electric 1.5 T system with a high-resolution antenna together with a homogenization phased array uniformity enhancement filter (fast spoiled gradient echo sequence, TR/TE/TI = 11.2/4.2/450 ms; flip angle 12°; slice thickness 1mm, 256 × 256 matrix and FOV 25 cm).

The source model was formed by placing a regular grid of 4560 sources spaced 1 cm apart forming a cube. Only sources corresponding to cortical regions according to the Anatomical Atlas Labelling were reconstructed and employed for further analyses, resulting in a total of 1210 brain sources. Subsequently, this source model was linearly transformed into each participant’s individual T1-weighted image. Single-shell head models were created using SPM12 brain segmentation.^34^ By combining head model and source model information, the lead field matrix was calculated using a modified spherical solution.

Sensor space signals were filtered between 2-45 Hz using a 450^th^-order finite input response filter designed with a Hann window. To avoid phase distortion, a two-pass filtering approach was used. In addition, to mitigate edge effects, 2000 samples of real data padding were added to each side of the signal. Time series of each source location were obtained using Linearly Constrained Minimum Variance beamformer as an inverse model. After source reconstruction, broadband signals were band-pass filtered into the five classical frequency bands, namely: delta (2-4 Hz), theta (4-8 Hz), alpha (8-12 Hz), beta (12-30 Hz) and gamma (30-45 Hz) for all subsequent analyses.

### Functional Connectivity and Graph Construction

Amplitude envelope correlation with leakage correction was employed to estimate functional connectivity between each pair of the 1210 brain sources, using the absolute value of the Pearson correlation between time-series envelopes as previously described.^35^ Typically, FC approaches in the literature have only considered connections between nodes in isolated frequency bands. Consequently, functional coupling is typically only estimated between nodes’ activities in the same frequency band, overlooking the relevance of cross-frequency couplings for various brain dynamics.^36^ In our approach amplitude envelope correlation was estimated for a given source so that its activity in each frequency band was correlated with that of every other source in all the frequency bands. This approach allows us to take into consideration not only the classical intra-band links (connection between two nodes in the same frequency band, e.g. *source 1 delta – source 2 delta* coupling), but also inter-band links (connection between two nodes in different frequency rhythms, e.g. *source 1 delta* – *source 2 alpha* coupling). By doing this, we obtained a set of 1210 (*sources*) by 5 (*bands*) connections for each node, resulting in a large and comprehensive connectivity matrix for each participant with 1210 by 1210 by 5 total connections (nodes by nodes by bands). These FC matrices were employed as the input for the graph analyses conducted in the present work.

A graph (*G*) is typically expressed as a set of nodes, denoted by the matrix (*V*); and the connections between those nodes, also called edges (*E*); *G = (V,E)*. The connections between nodes are commonly expressed in a weight matrix (*W*). In this matrix, the weight (*w*) of the connection between the node *i* and the node *j* is captured by the element w*_ij_* of the graph and represents the distance between two nodes in the graph.

In the context of a brain network study, the different nodes represent the regions of the brain and edges are usually expressed by a coupling or connection metric that can either be functional or structural. In our study, each of the 1210 sources was considered a node of the network. Furthermore, each frequency band was used as a different layer in our multilayer network, thus resulting in a total set of 6050 nodes (1210 nodes by 5 layers). A common problem in brain studies using network theory is the computational complexity of weighted networks which has typically led to the use of binarized networks. In this approach, an arbitrary threshold is employed to discard every connection below that value, maintaining only those above it. However, this methodology is not devoid of limitations, since the arbitrary nature of the threshold heavily influences the results obtained from the graph^37^, and a lack of clear consensus for how to establish an appropriate threshold hinders this approach. In our work we opted to use weighted undirected graphs in which the FC matrices previously described were employed as the weight of each edge in our graph.

### Centrality calculation

Centrality has been typically studied using different metrics and indicators, each of them capturing slightly different aspects of weight distribution in brain communication. In our work we decided to employ an integrative approach by combining three different network centrality metrics in a single measure that was employed for all the analyses. All the metrics were calculated for each network node of each subject using the NetworkX library (version 2.6.3) and SciPy library (version 1.7.1) in Python (version 3.10.4).

In particular, we used three graph measures as centrality indicators: node strength, eigenvector centrality and betweenness centrality. The strength of a node is determined by the number of connections a node has with other nodes of the network and the weight of its edges. It is calculated as the sum of the weights of the links incident to node *i*, where *w_ij_* is the weight of the link between nodes *i* and *j*.

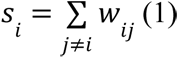

The eigenvector centrality considers not only the number and weight of connections of a node, but also the relevance of the nodes they are connecting to. In essence, a node is considered more central if it is connected to nodes that are also highly central themselves. It is calculated following *equation 2*, in which *M* is the adjacency matrix, λ the eigenvalues and *v* the eigenvectors. The element *i* of the eigenvector *v* that is associated with the largest eigenvalue λ_1_ holds the value for the eigenvector centrality of node *i*.:

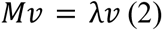

Finally, the betweenness centrality of a node estimates its ability to act as a bridge or intermediary between other nodes, by determining the proportion of shortest paths in the brain that runs through a given node. *Equation 3* describes betweenness centrality, where *V* is the set of nodes, σ(*j*, *k*) is the number of shortest paths, and σ(*j*, *k*|*i*) is the number of those paths passing through some node *i* other than *j*, *k*. As betweenness centrality takes link weights as a distance measure, we used the inverse of the weights (which, in our case, indicate functional proximity).

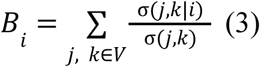

To reduce computation time, betweenness centrality was estimated using the algorithm developed by Brandes and Pich^38^ with the value of *k* set to 250.

This resulted in three vectors per subject, each of them with a length of 6050 (composed by 1210 values, one for each source, in each of the five frequency bands). To obtain a comprehensive measure of centrality, we combined the three metrics into one, hereafter referred to as *centrality score*. To do so, we first z-scored each of the metrics independently, considering their highly different numeric ranges. Afterwards, these vectors were averaged for each subject, obtaining a vector of the same length for each subject containing the centrality score employed in all the subsequent analyses.

### Statistical analysis

Statistical analyses were carried out in a stepwise manner. Firstly, we studied whether changes in brain network centrality were associated with the p-tau231 and NfL levels, individually, of each participant. To unveil the specific topographic distribution of said association we conducted a Spearman correlation analysis for each individual source in the brain, studying the statistical association between NfL/p-tau231 level with the centrality score of each individual brain source in each frequency band. To account for multiple comparisons, we conducted two-sided cluster-based permutation tests, using a Montecarlo approach.^39^ Significant sources were grouped based on their spatial contiguity, obtaining a cluster size defined as the sum of the individual statistics of each source in it. We conducted 10.000 permutations on the original data to create a surrogate distribution of random cluster sizes to compare our original cluster size to the surrogate size distribution. An alpha level of 0.05 was used to deem the original cluster as significant.

The second step of our analysis aimed to study whether associations between p-tau231 and centrality in each source were mediated by the grand-average centrality, (i.e. the absolute relevance of that source for brain communication). To do so, firstly we calculated the grand-average centrality of each source. This was done by simply averaging the centrality scores of each source for the five bands in a single centrality score for each brain source of each subject. This value was then averaged again across subjects to obtain a single centrality score value for each source for the whole population, thus resulting in a 1210 by 1 vector. These values reflect the specific relevance of each node for maintaining brain communication in the population. Finally, we studied the statistical association between the rho value obtained for each source in the first analysis step with the subsequently calculated grand-average of that source. This was done by calculating the correlation between the two mentioned vectors; namely, grand-average centrality (1210 values) and rho-value obtained in step one of the statistical analyses (1210 values).

## Acknowledgements

We would like to thank all the participants that have selflessly given us their time and made this study possible.

